# SSEalign: accurate function prediction of bacterial unannotated protein, based on effective training dataset

**DOI:** 10.1101/200915

**Authors:** Zhiyuan Yang, Stephen Kwok-Wing Tsui

## Abstract

The functions of numerous bacterial proteins remain unknown because of the variety of their sequences. The performances of existing prediction methods are highly weak toward these proteins, leading to the annotation of “hypothetical protein” deposited in NCBI database. Elucidating the functions of these unannotated proteins is an urgent task in computational biology. We report a method about secondary structure element alignment called SSEalign based on an effective training dataset extracting from 20 well-studied bacterial genomes. The experimentally validated same genes in different species were selected as training positives, while different genes in different species were selected as training negatives. Moreover, SSEalign used a set of well-defined basic alignment elements with the backtracking line search algorithm to derive the best parameters for accurate prediction. Experimental results showed that SSEalign achieved 91.2% test accuracy, better than existing prediction methods. SSEalign was subsequently applied to identify the functions of those unannotated proteins in the latest published minimal bacteria genome JCVI-syn3.0. Results indicated that At least 99 proteins out of 149 unannotated proteins in the JCVI-syn3.0 genome could be annotated by SSEalign. In conclusion, our method is effective for the identification of protein homology and the annotation of uncharacterized proteins in the genome.

## 1 Introduction

Because of the recent advance of DNA sequencing technology, abundant protein sequences are deposited in the NCBI RefSeq database and EBI UniProt database [1]. Unfortunately, the annotations remain unknown for a large amount of these sequences. Elucidating the function of the unannotated protein is an important topic of research in computational biology. It is a common task to annotate newly identified proteins by homology search in protein sequence databases of known other species. In molecular biology, homology is described as a relationship where two genes share a common ancestor, such as the relationship of human p53 gene and the mouse p53 gene. Homology can be divided into orthology and paralogy according to whether they are present in the same species. In this study, we mainly focused on the homology in different species, i.e. orthology.

Protein sequence comparison is the primary way for establishing homology. The routine method for annotation of protein-coding genes is identification of homologs by sequence alignment. For closely related proteins, homolog can easily be detected using conventional BLAST algorithm [2]. However, a remarkable challenge exists in the field because many evolutionarily distantly related proteins may vary highly at the amino acid level, especially for bacterial proteins. The failure of homology detection of a protein will lead to “hypothetical protein” or “uncharacterized protein” in its annotation.

In this study, we report a method that is specifically designed for homology identification of hypothetical proteins. Many tools are currently available for the detection of distantly related homology [3–5]. The two well-established tools are phmmer [4] and HHpred [5], both of which are widely used in homology identification in different species. Although these tools are supposed to be highly sensitive for remote homology detection, their performance in detecting protein homology in bacteria is sub-optimal. This problem is particularly important in the novel genome annotation, which relies on sequence alignment to annotate the function of translated proteins. The reason for their failures is that they have used datasets of protein homology with a wide range of identities but not datasets within bacteria, whose alignment identity of protein homology often fall into the twilight zone (≤35%) [6]. To overcome this shortcoming, we strictly restricted the training dataset to be protein homology in the bacteria and specifically optimized parameters for detecting homology in this situation.

In synthetic biology, one of the key focuses is how to build a minimal artificial cell which can provide suitable chassis for basic functional study. In 2010, the first version of minimal bacterial genome JCVI-syn1.0 had been reported by *Gibson* group with completely chemical synthesis, subsequently, this genome had been transplanted into the cell of *Mycoplasma capricolum* whose nucleus has been removed. This artificial genome had finally been demonstrated to possess the potential to self-replication and alive in the basic culture medium [7]. Later, transposon mutagenesis technique [8] was applied to the genes of JCVI-syn1.0 to identify dispensable genes. Finally, a more compressed genome JCVI-syn3.0, which is smaller than any genomes of autonomously replicating cells reported before, was obtained [9]. The JCVI-syn3.0 genome is approximately 531kbp in size and consists of 473 essential genes (438 protein-coding and 35 RNA-coding genes). The 438 protein-coding genes were then annotated by searching against TIGRfam database [10] and divided into two groups:

a. kno3.0 genes: 289 genes whose functions are clearly known, including those genes whose functions are extensively studied and can be supported by multiple aspects of the evidence, such as genomic context and the structure information.
b. unk3.0 genes: 149 genes whose functions are ambiguous and even completely unknown. As all these genes are essential for living organisms, we hypothesize that the 149 encoded proteins with unknown function should share homology with proteins in other bacteria.

Previous studies have suggested that the evolution rate of the protein secondary structure (SS) is much slower than that of the amino acid [11]. After the evolution of million years, the amino acid sequences have greatly changed but their structures have not been disrupted. The protein secondary structure is also the basis of the tertiary structure of a protein and it can serve as a bridge that links the primary sequence and the tertiary structure for the functional analysis.

The tools for protein secondary structure prediction have been extensively studied since the 1990s. These tools can be typically divided into two different categories: template-based prediction and *ab initio* prediction. Many of these computational methods are based on the close templates but a few for *ab initio* prediction. Among these tools, three of them (JPRED4 [12], PsiPred [13] and SSpro [14]) have been widely used for protein secondary structure prediction. JPRED4 is a web-based tool and the query sequence has a limit of protein length (≤800 amino acids), making it not suitable for SS prediction in this study. PsiPred is a template-based tool for secondary structure prediction. Considering that close templates could not be available for the protein homology in the twilight zone, PsiPred is also not suitable for SSEalign. Recently, a study reported a comparison of the prediction accuracy of these tools using third-party datasets. They found that SSpro has the best prediction accuracy for CullPDB and CB513 datasets [15]. Furthermore, the *ab initio* prediction method had been highly improved recently, which makes it possible to predict the secondary structure of those proteins in the twilight zone. Therefore, SSpro with *ab initio* strategy was employed to conduct secondary structure prediction in SSEalign.

In this study, we first trained our method in 20 well-studied bacteria to show the performance of the SSEalign. The line search optimization approach was used to derive the best scoring matrix for this application. The derived parameters were then applied to identify the homology among JCVI-syn3.0 genome and several other well-annotated bacteria. Lastly, six variables were used to evaluate our homology results.

## 2 Materials and Methods

### 2.1 Benchmark Dataset

The main objective of this work is to investigate how secondary structure information can be used to identify the corresponding homology for a set of protein sequences in bacteria. We need a set of protein sequences (benchmark dataset) with known homology information to train and to test our method. Thus, the protein-coding genes in following 20 well-studied bacteria [16] were selected as benchmark dataset: *Bacillus anthracis*, *Bacillus subtilis*, *Bifidobacterium longum*, *Clostridium botulinum*, *Clostridium tetani*, *Escherichia coli*, *Haemophilus influenzae*, *Helicobacter pylori*, *Lactobacillus acidophilus*, *Mycobacterium tuberculosis*, *Mycoplasma genitalium*, *Pseudomonas aeruginosa*, *Rickettsia prowazekii*, *Salmonella typhimurium*, *Staphylococcus aureus*, *Streptococcus pneumoniae*, *Streptococcus thermophilus*, *Thermotoga maritima*, *Vibrio cholerae*, *Yersinia pestis*. Genome annotation files of these 20 bacteria were obtained in Ensembl database [17].

The benchmark dataset consisted overall positive and negative samples. The positive samples were composed of all homologous protein pairs in different species, for example, protein rpiL in *E. coli* and *B. subtilis.* After applying this criterion, 75,206 protein pairs were deposited into the positive dataset. If we select all the protein pairs as negative controls, the number of samples will be extremely large, leading to an extreme imbalance when compared with the positive dataset. Such imbalance will in turn dramatically affect the performance evaluation of model training in this type of machine learning problem [18, 19]. Therefore, only 75,206 non-homologous protein pairs were randomly picked to constitute negative dataset. To validate the robustness of our method, we repeatedly conducted this procedure of random sampling for 100 times to obtain different negative dataset for the downstream analysis.

In literature, the benchmark dataset usually divided into a training and a testing dataset: the former is for the training the model, while the latter is for testing it. We randomly divided the benchmark dataset into training and test datasets using the ratio of 9:1 (10-fold cross-validation). Thus, the training dataset consisted of 67685 homologous (TRN-POS) and 67685 non-homologous (TRN-NEG) pairs, which the testing dataset consisted of 7521 homologous (TST-POS) and 7521 non-homologous (TST-NEG) pairs.

### 2.2 The <u>B</u>asic <u>A</u>lignment <u>E</u>lements (BAEs) in SSEalign

After obtaining the training dataset, the SSpro toolkits were used to conduct the protein secondary structure prediction using the *ab initio* prediction model with default parameters. The training dataset containing information of secondary structure element (SSE) was then created (Figure 1). To better show the actual performance of our method, the sequence with simple repeats were excluded. It has long been recognized that three different types of protein secondary structure are present in nature. In this study, the one-character alphabet was used to represent these secondary structures:

*H* = Helix (mainly alpha helix);
*E* = Sheet (mainly beta sheet);
*C* = Random coil;

**Figure 1.**
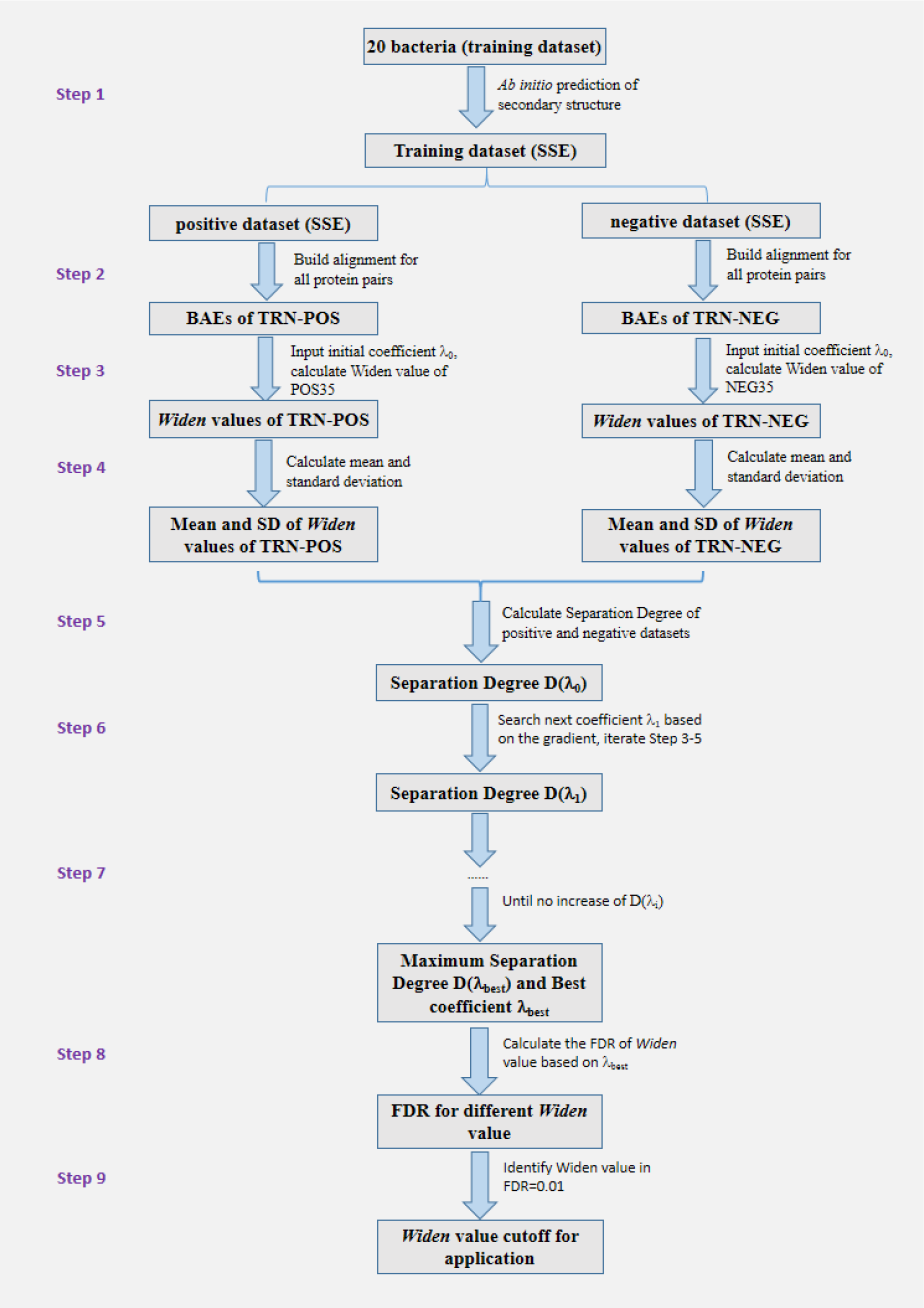
The flowchart of the SSEalign pipeline. SSE: secondary structure element; TRN-POS: positive samples in training dataset; TRN-NEG: negative samples in training dataset; BAEs: basic alignment elements, i.e. [HH, EE, CC, HE, HC, EC, GN, GO]; SD: Standard Deviation.

To better evaluate the performance of our method, sequences with simple repeats were excluded. The EMBOSS-stretcher tool [20] was used to conduct the global sequence alignment of the SSEs of protein pairs. EMBOSS (European Molecular Biology Open Software Suite) is a free open source software for molecular biology. Within the EMBOSS, the stretcher tool is an effective package for global alignment. After finishing this process, the alignments were segregated into eight basic alignment elements.

Eight possible BAEs could be found in the secondary structure alignment generated by SSEalign:

HH = *H* coordinates with *H* in the SSEalign
EE = *E* coordinates with *E* in the SSEalign
CC = *C* coordinates with *C* in the SSEalign
HE = *H* coordinates with *E*, or *E* coordinates with *H* in the SSEalign
HC = *H* coordinates with *C*, or *C* coordinates with *H* in the SSEalign
EC = *E* coordinates with *C*, or *C* coordinates with *E* in the SSEalign
*GN* = number of gaps in the SSEalign
*GO* = number of gap openings in the SSEalign

### 2.3 Scoring system of SSEalign

It is very probable that the contribution of eight BAEs was very different in the sequence alignment. To optimize the performance of the SSEalign, we developed a customized evaluation index: *Widen* (<u>W</u>eighted <u>ident</u>ity), to evaluate the alignment. The *Widen* value varies in the range [0%, 100%] and it can be recognized as the analog of “alignment score” generated by BLAST. Then, similarity of a two-sequence alignment can be described by the index *Widen*: *Widen* = 0%, totally different; *Widen* = 100%, completely identical. The formula of *Widen* is shown in formula (1)

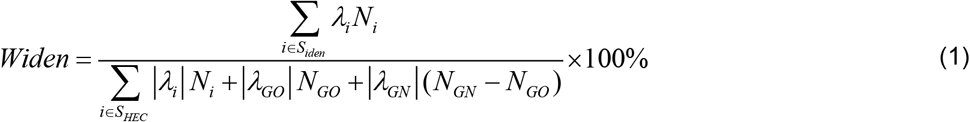

*S*_*HEC*_: is the set of all the possible BAEs consisted of three types of secondary structures in the sequence alignment, i.e. *S*_*HEC*_ = {*HH*, *EE*, *CC*, *HE*, *HC*, *EC*}.
*S_iden_*: is the set of all the identical matches of BAEs, i.e. *S_iden_* = {*HH*,*EE*, *CC*}
*λ*_*i*_: is the coefficient of BAEs, i.e. λ = [*λ*_*HH*_, *λ*_*EE*_, *λ*_*CC*_, *λ*_*HE*_, *λ*_*HC*_, *λ*_*EC*_]
*N*_*i*_: is the number of correspondent BAEs in the alignment.
*λ*_*GN*_: is the coefficient of gap, i.e. penalty for gap extension.
*N*_*GN*_: is the number of gaps in the alignments.
*λ*_*GO*_: is the coefficient of gap openings, i.e. penalty for opening the first gap in the alignment
*N*_*GO*_: is the number of gap openings excluding those gaps at both ends.

### 2.4 Optimization method in training dataset

In this optimization process, we try to minimize the overlap of *Widen* values of positive and negative dataset: The *Widen* values of the positive dataset could be as large as possible meanwhile the *Widen* values of negative dataset could be as small as possible. Thus, we defined the following formula (2) to indicate the separation degree of the positive and negative dataset:

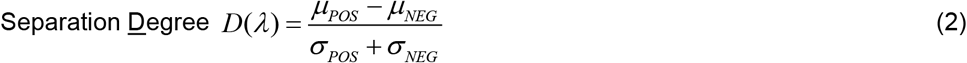

*μ*_*POS*_: is the mean of *Widen* values in the positive dataset.
*μ*_*NEG*_: is the mean of *Widen* value of values in the negative dataset.
*σ*_*POS*_: is the standard deviation of *Widen* values in the positive dataset.
*σ*_*NEG*_: is the standard deviation of *Widen* values in the negative dataset.

To maximize the separation degree in our study, the backtracking line search approach was applied to derive the best scoring coefficient *λ* to separate the positive and negative dataset. The backtracking line search approach is an efficient iterative method to find extreme points based on a start point. In this optimization process, we used the following *λ*_0_ as a start point to derive the best scoring matrix.

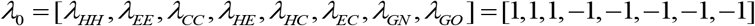

### 2.5 Criterion for homology identification

False discovery rate (FDR) was introduced to identify the appropriate threshold of *Widen* value for homology identification. The FDR is defined as the expected ratio of false positives among the predicted significant results and serves as a more convincing scale than p-value scale because of its directly useful interpretation [21]. The FDR will be 0.5 if identifying homology by random selection FDR value was defined in the following formula (3).

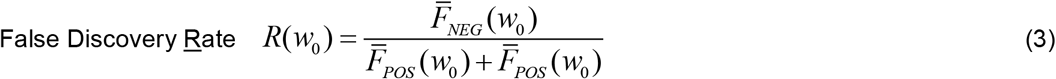

*w*_*o*_: is the given *Widen* value.
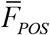: is the complementary cumulative distribution function of *Widen* values of the positive dataset.
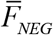: is the complementary cumulative distribution function of *Widen* values of the negative dataset.

The FDR values towards different *Widen* values were calculated. The criterion FDR=0.01 was frequently used as a cutoff in previous biological studies [22–24]. Thus, this cutoff was also adopted in the study for the homology identification.

### 2.6 The performance of SSEalign and compared methods on testing dataset

Currently, several tools were published for distantly related homology identification, such as phmmer and HHpred. The phmmer is a toolkit of HMMER3 which detect the homology via sequence profile comparison while the HHpred is a toolkit of HHsuite which detect the homology by HMM-HMM comparison. These tools were benchmarked with our SSEalign tool based on testing datasets. For phmmer, a cutoff e-value≤1e-5 was used in the all-against-all strategy in the testing dataset. For HHpred, each protein was searching against the testing dataset with HHBlits as the HMM generation method.

To illustrate the performance of SSEalign and compared methods, the ROC (Receiver Operating Characteristic) curves were plotted. The ROC curve plots true positive rate (*Sensitivity*) against the false positive rate (1-*Specficity*) and their definitions were shown in formulas (4) and (5)

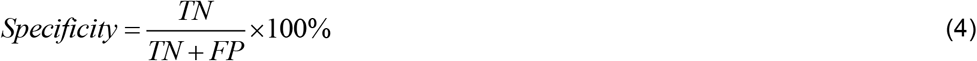

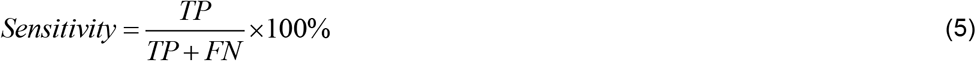

In the formulas, TP: true positives; FN: false negatives; TN: true negatives; and FP: false positives.

The best prediction model could produce a point with the coordinate (0,1), which indicates 100% true positive rate and 0% false positive rate. To avoid the threshold setting bias, the area under the curve (AUC) was frequently used to show the performance quality of binary classification methods [25,26]. A completely random guess will yield the AUC value 0.5, while a perfect model will yield the AUC value 1.0.

### 2.7 SSEalign Application: JCVI-syn3.0 genome annotation

Very recently, *Gibson* group has published the latest minimal bacterial genome JCVI-syn3.0 which was only consisted of 473 genes (438 protein-coding genes and 35 RNA-coding genes). The 438 protein-coding genes can be divided into 289 proteins with clear function (kno3.0 dataset) and 149 proteins with unknown functions (unk3.0 dataset).

The method of pairwise BLAST with the cutoff E-value≤1e-5 was applied in the kno3.0 dataset to identify the homology among the genomes of JCVI-syn3.0 and the 20 selected bacteria. The numbers of shared genes in the 20 bacteria were further enumerated. The top three species (*E. coli*, *B. subtilis and H. influenzae*) sharing the highest number of genes with JCVI-syn3.0 were selected for SSEalign analysis. The secondary structures of unk3.0 protein and proteins in these three selected bacteria were then predicted by the toolkit SSpro. The pairwise global sequence alignments of the generated secondary structure elements were conducted by EMBOSS-stretcher with the scoring matrix derived from the process of “Optimization Method”. The hitting sequences with FDR ≤ 0.01 were designated as homologous protein candidates.

To evaluate the prediction accuracy of our method, the following six parameters of homologous protein candidates were applied.

(a) Protein domain: We hypothesize that homologous proteins tend to share the same protein domains. The domain information of these proteins was predicted by Pfam [27] and InterProScan [13] by using the default parameters. If the protein pairs of homology candidates share the same domains, then the two homologous protein candidates were considered supported in the parameter of protein domain.

(b) Feature binding site: We hypothesize that feature binding sites, such as ATP-binding site and metal binding site, are conserved in homologous proteins during evolution. The homologs of unk3.0 proteins in *E. coli, B. subtilis* and *H. influenzae* were singled out to conduct the multiple sequence alignments by CLUSATLX using default parameters. The feature binding sites of these proteins in the three species were then retrieved from the UniProt database. If the corresponding featured binding sites could be found in unk3.0 proteins, the homologous protein pairs were considered supported in the parameter of feature binding sites.

(c) Gene synteny: We hypothesize that homologs tend to share the same gene synteny in different genomes, which means the upstream and downstream genes were the same for a certain gene in different species [28]. The gene loci of proteins in each genome were identified by searching against the corresponding genome with tBLASTn. It has been reported that average length of synteny block among distant species is about 150kbp [29]. Thus, the neighboring genes within 75kbp upstream and 75kbp downstream of a certain gene were compared in each species. The homologous protein pairs with the same corresponding gene synteny were considered supported in the parameter of gene synteny.

(d) Protein-protein interaction: As these genes were known as minimal essential genes, we hypothesize that their encoded proteins tend to interact with each other [30]. The protein-protein interaction (PPI) datasets of *E. coli* were obtained from BioGrid database, which includes PPI network of *E. coli* genes curated from most recently published papers [31]. The PPI network of homologous proteins of JCVI-syn3.0 predicted by BLAST (kno3.0) and SSEalign (unk3.0) in *E. coli* were then constructed by Cytoscape [24]. If the homologous proteins predicted by SSEalign in *E. coli* can interact with those predicted by BLAST, the homologous protein pairs were considered supported in the parameter of protein-protein interaction.

(e) Essential gene: We hypothesize that the genes in JCVI-syn3.0 are also essential genes in other species. We collected 296, 271 and 431 essential genes of *E. coli*, *B. subtilis* and *H. influenzae, respectively*, from previous studies [32–34]. If the homologous genes were also essential genes in *E. coli, B. subtilis* or *H. influenzae*, these homologous gene pairs were considered supported in the parameter of essential gene.

(f) Phylogenetic topology: We hypothesize that homologous proteins tend to cluster with each other in the phylogenetic tree [35]. The unk3.0 proteins and their homologs in other bacteria were selected to construct phylogenetic trees by CLUSTALX and MEGA [36]. If the unk3.0 protein and its homologs can be clustered in a single branch in the phylogenetic tree, the homologous protein pair was considered supported in the parameter of phylogenetic tree topology.

The rigorous criteria for these parameters were applied to the homologous protein candidates to check if they were supported in each parameter. The number of supported parameters for each homologous protein pair was then calculated and the cumulative distribution was plotted. The homologous protein candidates which have support in at least three parameters were considered as homologs in this study.

## 3 Results and Discussion

### 3.1 The performance of SSEalign and compared method

After screening proteins within 20 well-studied bacteria, 75,206 homologous protein pairs (positive samples) and 75,206 randomly selected non-homologous protein pairs (negative samples) were selected as the benchmark dataset. The benchmark dataset was divided into training datasets (67,685 homologous and 67,685 non-homologous pairs) and testing datasets (7,521 homologous and 7,521 non-homologous pairs) by the 10-fold cross-validation. The secondary structure elements (SSEs) of training dataset were then predicted by the SSpro toolkits to get the Training-SSE dataset.

Subsequently, the global alignment was applied to the protein pairs in positive and negative datasets. The numbers of basic alignment elements (BAEs) of each alignment were then recorded and transformed into the alignment information matrix. This alignment information matrix and the initial values of scoring matrix *λ*_0_ = [1,1,1,–1,–1,–1,–1,–1] were inputted into the backtracking line
search script in MATLAB software to derive the best scoring matrix for SSEalign. After many thousands of iterations, we obtained the best scoring matrix for each BAE of the global alignment. We found that the prediction accuracy was not affected by the random selection of negative dataset, indicating that the SSEalign is robust in the different negative datasets.

The best scoring matrix for each BAE of the global alignment were showed in following equation:

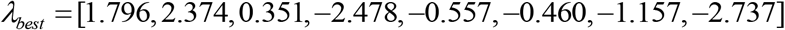

In this equation, the indicated values were the correspondent coefficients of [*HH*, *EE*, *CC*, *HE*, *HC*, *EC*, *GN*, *GO*], respectively. As expected, the helix (*H*) and sheet (*E*) are more indicative in sequence than the random coil (*C*), with identity coefficients of 1.796 and 2.374, respectively. Thus, the penalty scores of “*HC*” and “*EC*” are obviously smaller than that of “*HE”*. A “*GO*” penalty score of −2.737 and “*GN*” score of −1.157 indicate that sequences differ greatly in length will not be homologous because abundant gaps will be present in the global alignment. Such gaps will significantly reduce the *Widen* value and alignments containing large numbers of gaps will be neglected during the homology identification.

Figure 2 shows the FDR of homology identification by different *Widen* values. Based on these results, the *Widen* value of 59.18% when FDR is 0.01 was identified. This indicated that the false possibility to identify two homologous sequences in the region of *Widen*≥59.18%, was lower than 1%. This *Widen* value was used as the significance cutoff in subsequent analysis.

**Figure 2.**
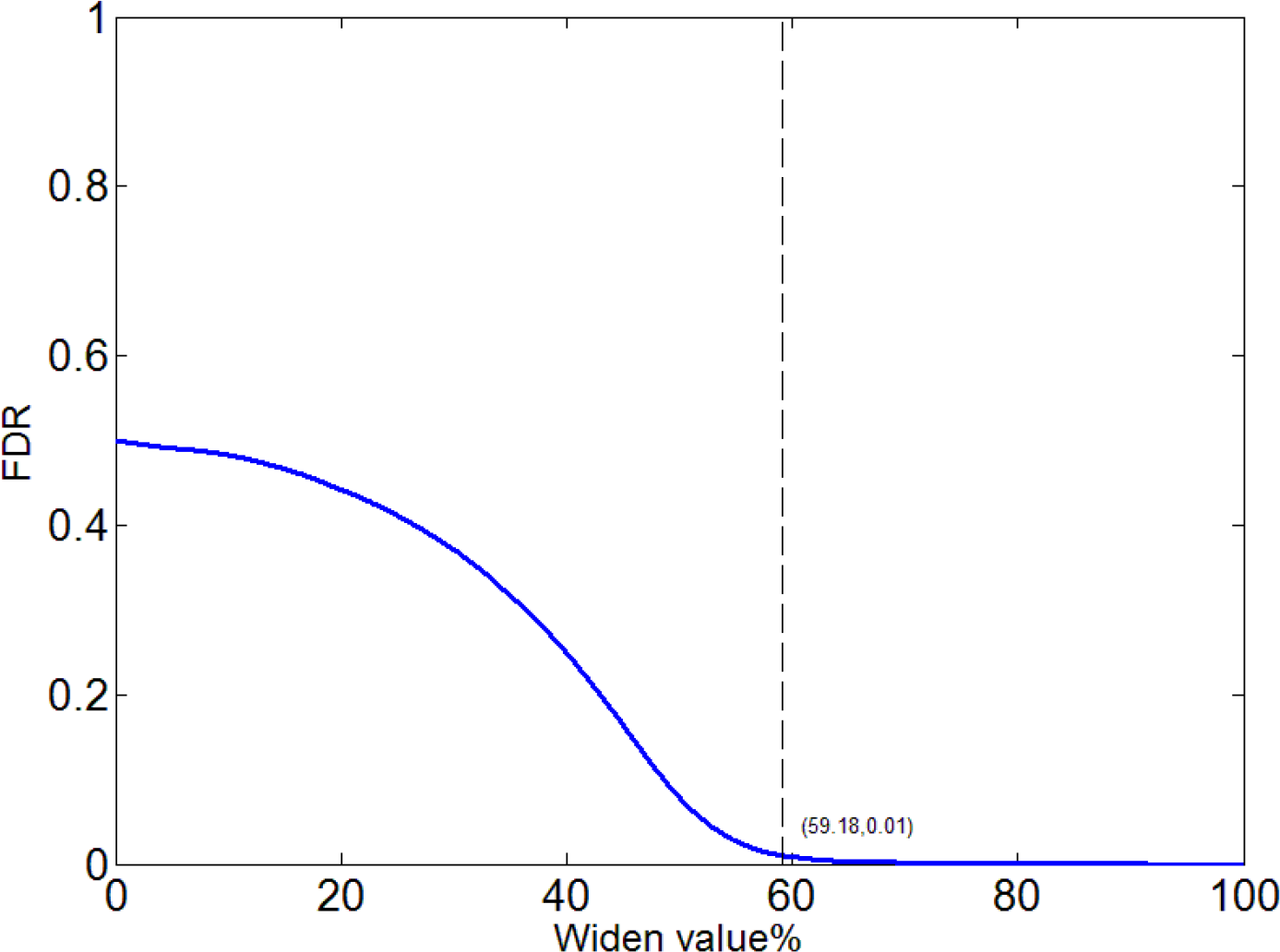
The FDR of homology identification by different Widen value. The dashed line shows the *Widen* value to achieve FDR=0.01

The comparison of performance of SSEalign and other available tools for homology identification based on the testing dataset were shown in Figure 3. For this dataset, the AUC values for three tools (SSEalign, HHpred and phmmer) were 0.912, 0.841 and 0.804, respectively. The SSEalign has an obvious better performance when compared with HHpred and phmmer, implying that our method is robust for homology identification in different species. The prominent performance of SSEalign will help us to re-annotate those proteins with unknown functions, especially for the bacterial proteins because their diversities were much higher than proteins of higher organisms.

**Figure 3.**
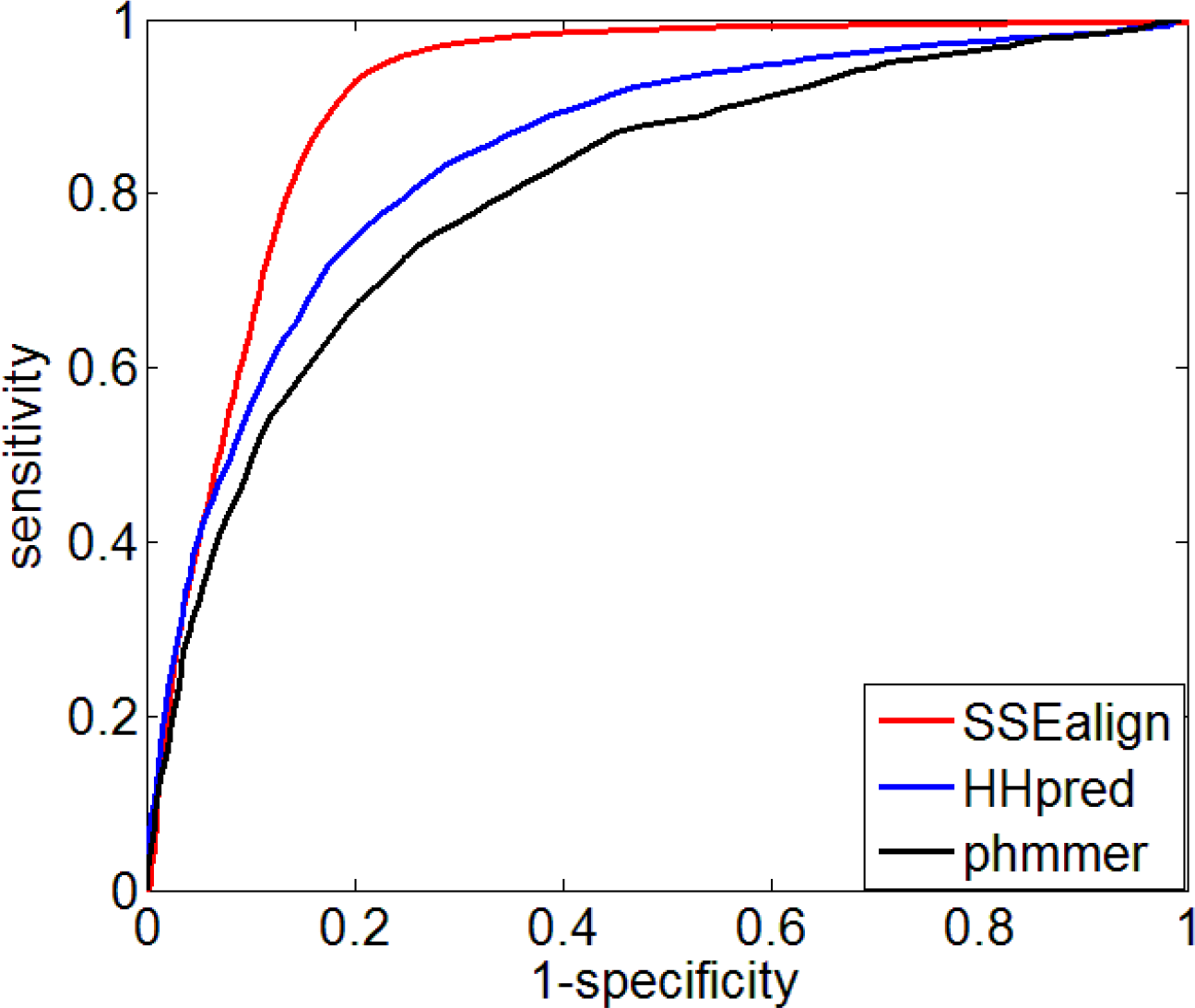
performance of SSEalign and compared method on the testing dataset. The red, blue and black curves indicate the performance of SSEalign, HHpred and phmmer, respectively.

### 3.3 Annotation of the genome JCVI-syn3.0

The genome JCVI-syn3.0 consisted of 438 protein-coding genes and 35 RNA-coding genes. The protein-coding genes could further be divided into kno3.0 dataset whose function was known and unk3.0 dataset whose function was unknown. A total of 289 proteins were present in the kno3.0 dataset and their identities could be easily identified by BLAST with a cutoff e-value≤1e-5. We found that the kno3.0 proteins shared 268 homologs with predicted proteins of *E. coli.* The numbers of homologs of the kno3.0 dataset in *B. subtilis* and *H. influenzae* were also very high (243 and 231, respectively).

Clustering of these 289 proteins showed that the top categories were 50S ribosomal proteins, 30S ribosomal proteins and DNA polymerases, which constituted 11.2% (30/289), 7.4% (20/289) and 3.3% (9/289), respectively (Table 1). The 50S and 30S ribosomal proteins are the basic components of prokaryotic ribosomes and they are highly conserved among all the species. Thus, these proteins can be easily annotated by conventional methods.

**Table 1.**
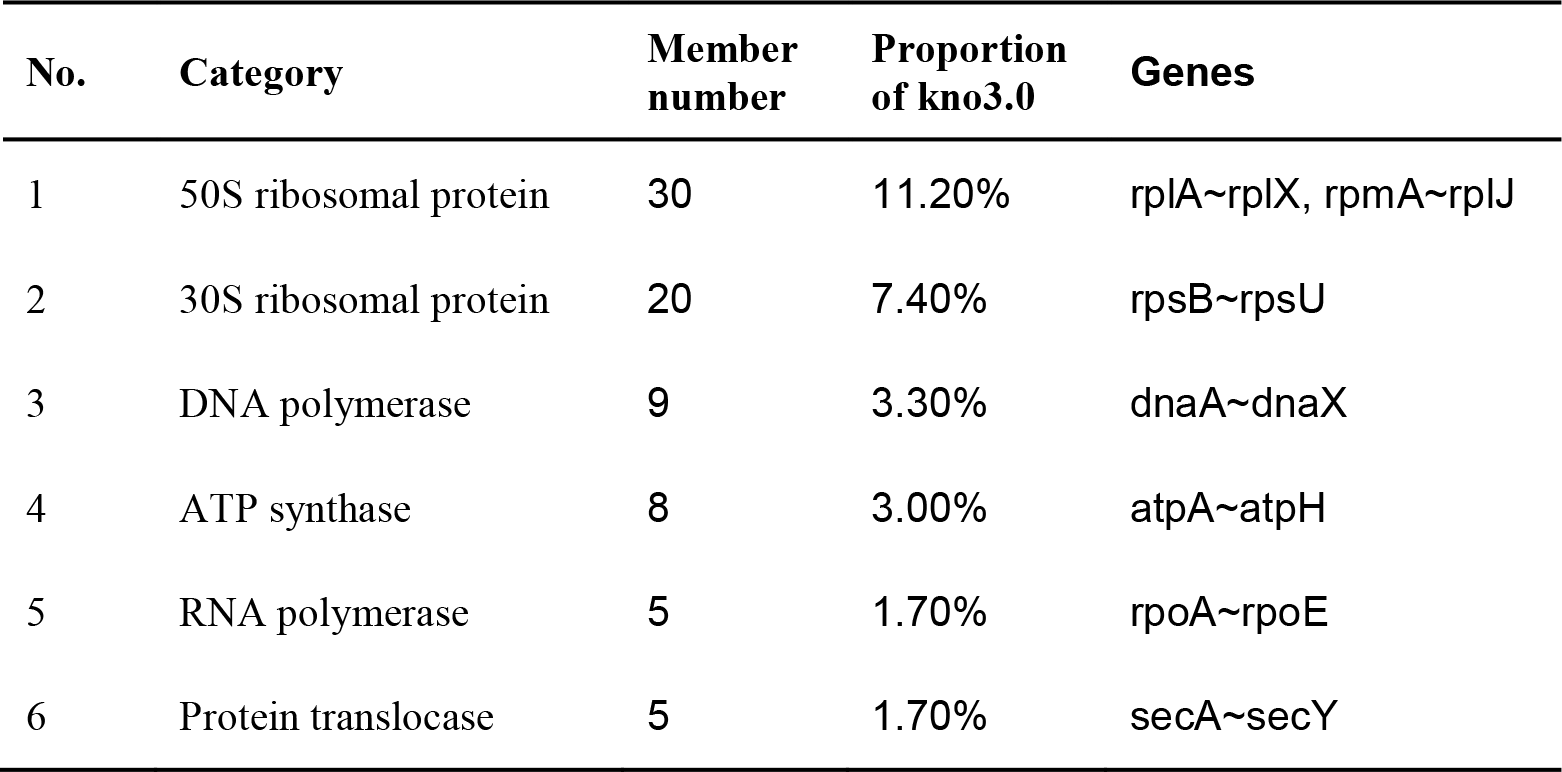
The main categories of identified proteins of kno3.0 dataset

We then identified homologs of the unk3.0 proteins in three bacteria (*E. coli*, *B. subtilis* and *H. influenzae)* using the SSEalign. Homologs with an FDR cutoff≤0.01 were selected as homology candidates for further evaluation. The performance of SSEalign is satisfactory because homology candidates could be found in 91.3% (136/149) of the unk3.0 proteins. We then assigned these 136 proteins into different functional categories by DAVID enrichment analysis and the results were shown in Table 2. The two largest groups of proteins were nucleotide-binding proteins (45 members, 30.2% of unk3.0) and ATP-binding proteins (43 members, 28.9% of unk3.0). The nucleotide-binding was a critical step for gene replication and transcription while ATP-binding was the determinant process for energy production and utilization in metabolic processes. Thus, it is not surprising that these proteins are essential for living cells. A relatively small group of proteins was transferases (40 proteins, 26.8% of unk3.0), which were commonly used for transferring the acetyl, methyl and phosphate group during the cell cycle, making them indispensable in the minimal bacterial genome It is well known that transferases and proteins for binding were highly diverse in different species [37], leading to the failure in previous homology identification because they fell into the twilight zone when searching against the TIGRfam database. However, the secondary structures of these proteins were highly conserved in all species, explaining why SSEalign has such an excellent performance for the annotation of these proteins.

**Table 2.** The main categories of identified proteins of unk3.0 dataset

| No. | Group name | Member number | Proportion of unk3.0 | Enrichment p-value |
| --- | --- | --- | --- | --- |
| 1 | Nucleotide-binding protein | 45 | 30.2% | 2.1E-14 |
| 2 | ATP-binding protein | 43 | 28.9% | 3.6E-14 |
| 3 | Transferase | 40 | 26.8% | 9.5E-07 |
| 4 | Hydrolase | 26 | 17.4% | 9.8E-06 |
| 5 | Transport protein | 21 | 14.1% | 2.8E-05 |
| 6 | Metal-binding | 20 | 13.4% | 2.2E-06 |
| 7 | Lyase | 19 | 12.8% | 1.6E-07 |
| 8 | Ligase | 18 | 12.1% | 7.6E-09 |
| 9 | Nucleotidyltransferase | 12 | 8.1% | 8.4E-07 |
| 10 | Helicase | 11 | 7.4% | 6.3E-09 |

Subsequently, the identified homologs were further evaluated using six parameters (protein domain, feature binding site, gene synteny, protein interaction, essential gene and phylogenetic topology) to validate the reliability of our method. Such parameters are independent functional supports of homology pairs identified by SSEalign. We found that 94.1% (128/136) of homologous proteins were supported by at least two parameters and 72.8% (99/136) of them were even supported by three or more parameters (Figure 4). Table 3 showed the identified homologs of the unk3.0 dataset with the top 10 *Widen* value in *E. coli*. Among these proteins, most of them were supported by at least four parameters, including MMSYN_0371 (annotated as cydC) and MMSYN1_0039 (annotated as ftsH).

**Figure 4.**
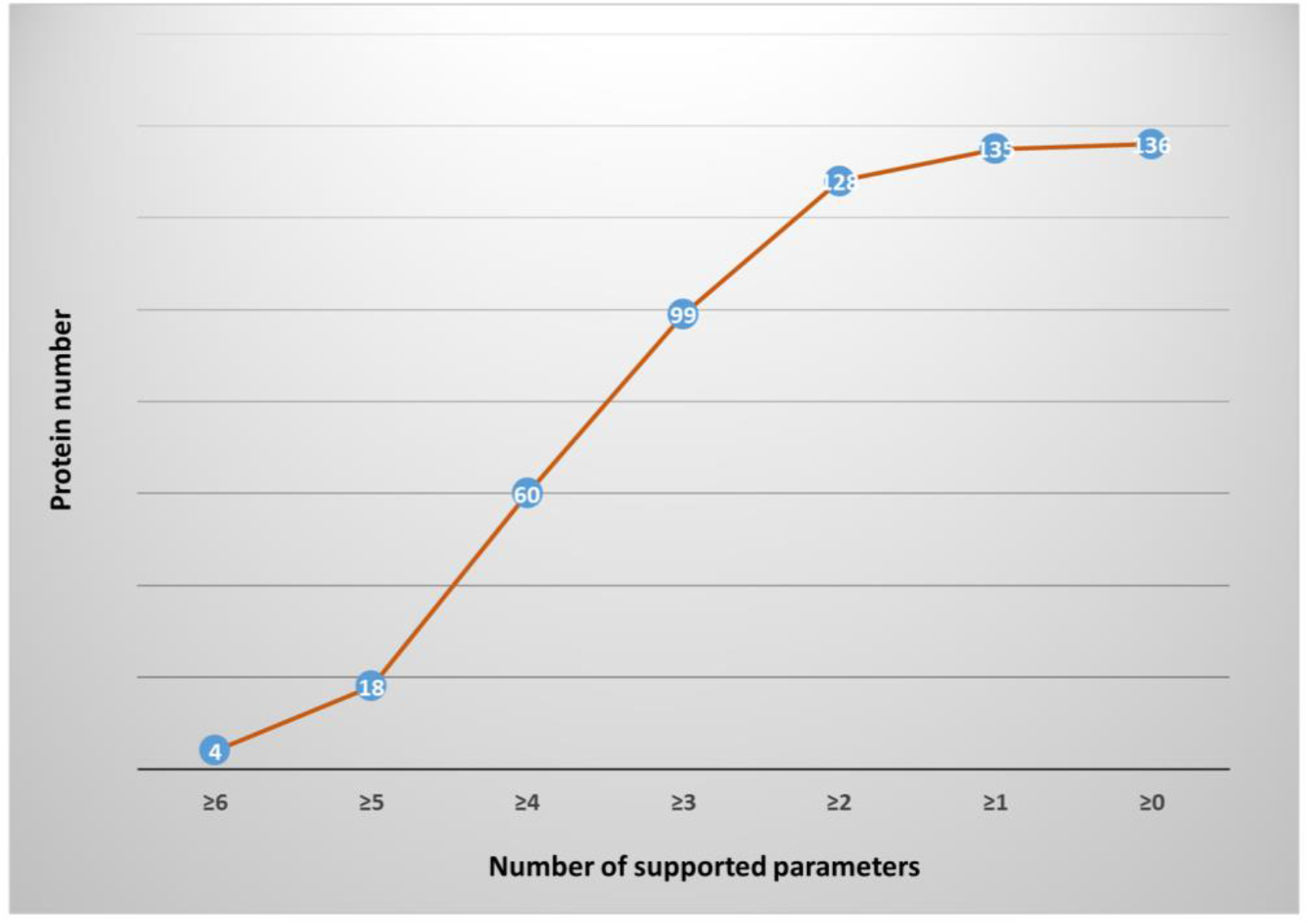
The numbers of proteins supported by different parameters. The x-axis indicated the number of six parameters: protein domain, feature binding site, gene synteny, protein-protein interaction, essential gene and phylogenetic topology. The y-axis indicates the number of proteins supported by the number of parameters in the correspondent x-axis.

**Table 3.**
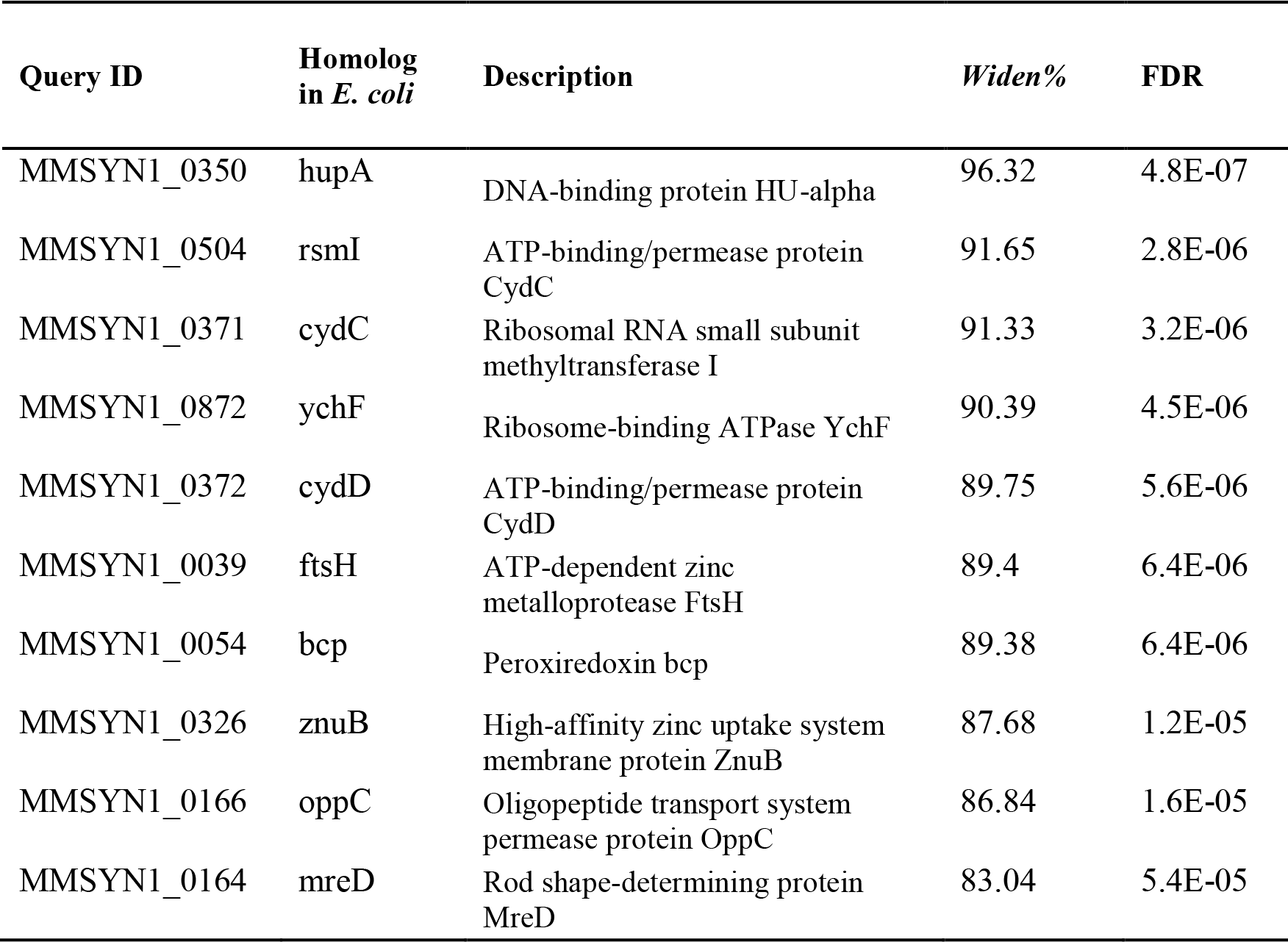
The top 10 identified protein of unk3.0 protein in *E.coli* genome. The p-value indicated that the possibility to achieve a correspondent *Widen* value by random selection. The FDR value indicated that the expected false discovery rate of claimed homology of these two proteins

The cydC and ftsH proteins are two kinds of highly diverse proteins in bacteria and the homology identifications of these two proteins in the new species by previous published computational biology method is very challenging [38, 39]. But their secondary structure is extremely conserved so our method can successfully detect it in JCVI-syn3.0.

The MMSYN1_0371 and MMSYN1_0039 shared extremely low FDR values, 3.2E-6 and 6.4E-6, with *E. coli* proteins cydC and ftsH, respectively. The multiple sequence alignment of ftsH proteins in 4 genomes (JCVI-syn3.0, *E. coli, B. subtilis* and *H. influenzae*) showed that 191 amino acids were exactly conserved. For the gene synteny analysis of *ftsH*, the *gyrA* is in its adjacent upstream region and the *lysS* is in its adjacent downstream region, which is consistent with their loci in the *B. subtilis* (Figure 5). In summary, these evaluation results further suggested that our annotation results of unk3.0 proteins by SSEalign were very convincing.

**Figure 5.**
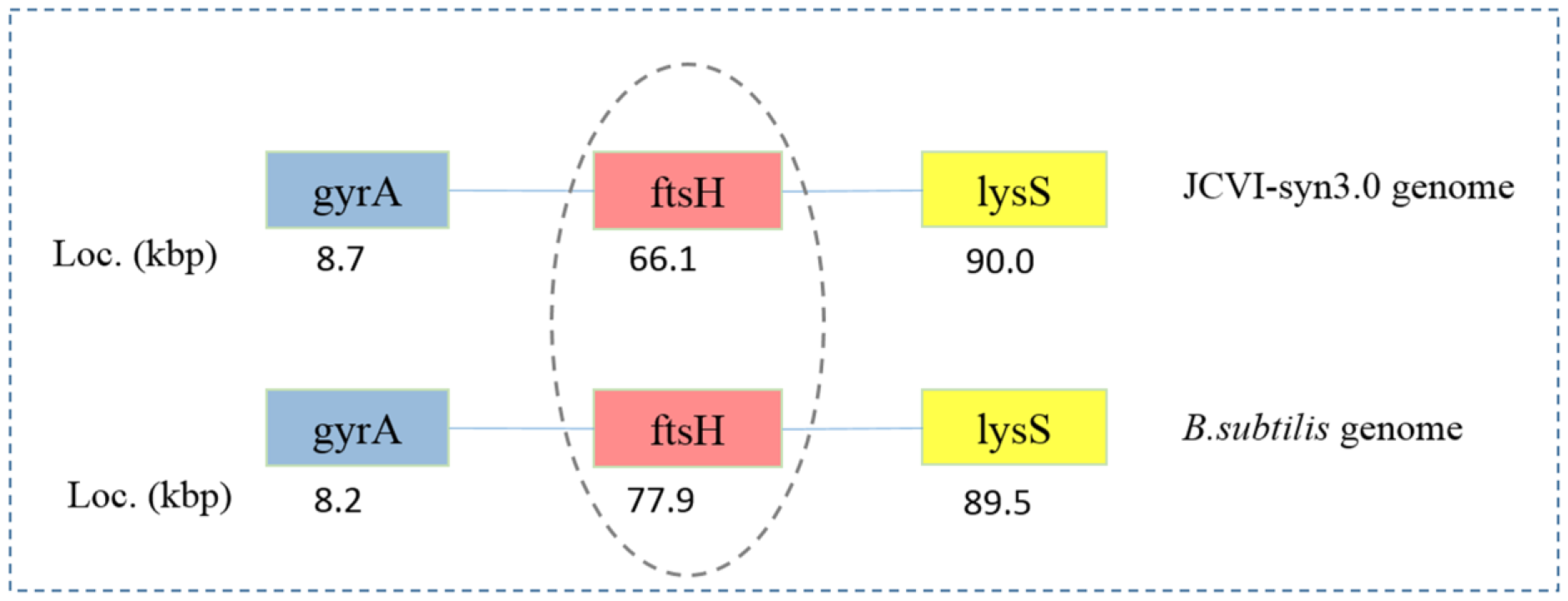
Gene synteny analysis of *ftsH* in genomes of JCVI-syn3.0 and *B.subtilis.* The numbers are the correspondent gene loci of in the genome. For the *ftsH*, its upstream and downstream genes are *gyrA* and *lysS*, respectively, in both JCVI-syn3.0 and *B. subtilis* genome

## 4 Conclusion

In conclusion, we developed a novel method to identify protein homology in the bacteria. The optimization method of backtracking line search was applied to obtain the best scoring matrix for secondary structure element alignment. Performance results on testing dataset showed that the SSEalign achieved a ROC value as high as 0.912, obviously better than existing prediction methods. The SSEalign was then applied to the minimal bacterial genome JCVI-syn3.0 to identify homologs of proteins that cannot be annotated using previous methods. Among these proteins, 99 members of them were considered homologous between the JCVI-syn3.0 genome and other well-studied bacteria. These genes have not been annotated in this genome before and may reveal new information about the essential mechanisms in living organisms. We have the confidence that the SSEalign and the evaluation strategy reported in this study are also useful for re-annotation of those proteins with the annotation of “hypothetical proteins” or “uncharacterized proteins” in the NCBI RefSeq database or EBI UniProt database. In conclusion, our method can remarkably fill the gaps in genome biology and expand the territory of systems biology.

## Acknowledgements

The authors are grateful to the supports and help of all labmates.

